# Novel female reproductive organ differentiates postmating transcriptional response to insemination versus arrival of sperm in bedbugs

**DOI:** 10.64898/2026.03.17.707905

**Authors:** Birte M. Martens, Caitlin E. McDonough-Goldstein, Oliver Otti, Susanne Broschk, Lucas Kullmann, Klaus Reinhardt, Martin D. Garlovsky

## Abstract

Following the evolution of internal fertilisation, the female reproductive tract became the site of reproductive interactions. However, our understanding of the evolution of female reproductive tract function, including postmating responses critical for reproductive success, are taxonomically limited. Traumatic insemination in the common bedbug (*Cimex lectularius*) presents an unusual scenario under which postmating responses unfold. Bedbugs have evolved a novel organ, the mesospermalege, that is the site of initial ejaculate × female interactions. As the female reproductive tract does not take receipt of the ejaculate until several hours after mating, bedbugs provide a unique opportunity to explore the evolution of a novel reproductive organ that decouples postmating female responses involved in mating and transfer of the ejaculate from sperm storage, ovulation, and oviposition. Here we show that the mesospermalege has a gene expression profile consistent with functions of ejaculate processing and immune response normally found in the lower reproductive tract of other insect species. In parallel, the postmating response in the lower female reproductive tract is delayed, coinciding with movement of sperm through the female, clearly showing that the postmating response has evolved in response to sperm receipt rather than being an innate function of the tissue. Notably, we also found expression of male seminal fluid genes in the mesospermalege, indicating that intersexual molecular dynamics influence the evolution of reproductive tissues. Our results provide insights into the evolution of novel reproductive traits and female postmating physiology in a global pest with an unusual reproductive biology.

**SIGNIFICANCE:** Reproduction poses one of the most persistent challenges faced by animals whereby females undergo a series of physiological changes after mating. The independent origin of a reproductive organ in bedbugs (called the mesospermalege) which has evolved to alleviate the costs of traumatic insemination presents a unique case to study the evolution of a novel trait and postmating physiology. Using transcriptomics, we show that many genes normally expressed in the female reproductive tract are instead expressed in the mesospermalege. The reproductive tract also shows a delayed postmating transcriptional response coinciding with sperm entry into the reproductive tract. Our results provide insights into the evolution of reproductive traits and female postmating physiology in a global pest with an unusual reproductive biology.

## INTRODUCTION

The evolution of internal fertilisation in animals introduced unique challenges in the process of sexual reproduction. Following this transition in reproductive mode, the female body shapes the physical and chemical environment that sperm must survive and traverse before fertilisation (1–3). In turn, mating can physically harm the female (4), introduce pathogens from the environment or in the ejaculate (5), and transferred seminal fluid proteins alter female physiology in ways which may be costly to female fitness (6). The outcome of these interactions between the sexes entails both cooperation and conflict, shaping the evolution of morphological and physiological reproductive traits (7–9).

There has been a growing appreciation for the role that females play in mediating postmating interactions and the evolution of reproductive traits in both sexes (10–16). For instance, female reproductive tissues and secretions support sperm viability after insemination (17–20); from hours or days in some mammals (21); to months or even years in some eusocial Hymenoptera (22). However, conflict between the sexes also shapes postmating physiological traits (2, 23, 24). In the cabbage white butterfly (*Pieris rapae*), females break down the hard spermatophore envelope with teeth-like signum in the bursa and have co-opted digestive enzymes from the gut to process the ejaculate (25, 26). The combination of selection pressures arising from, or constrained by, the multiple intersecting roles of the female reproductive system thus differentially influence the evolution of these tissues across species. These functions include (i) mitigate the costs of mating, including infection (27), (ii) process the ejaculate, (iii) aid and potentially modulate sperm transport, storage, and survival (11, 18, 28), and (iv) prepare for egg production, maturation, and ovulation (29–32).

To accommodate these various functions, females of many species undergo a series of changes in morphology, physiology and behaviour after mating (33–35). These postmating responses are accompanied by dynamic changes in gene expression in the female reproductive tract (12, 13, 34–41). For instance, the immune challenge associated with mating coincides with an increase in immune gene expression in the female reproductive tract, a pattern found from insects to mammals (42–48). In *Drosophila melanogaster*, the earliest transcriptional changes in the reproductive tract occur within minutes after courtship and mating, with a peak in differential gene expression 6 hours after the start of mating (12, 38, 49). The female reproductive tract is hypothesised to start in a “poised” state and after mating undergo terminal differentiation to support subsequent reproductive events (e.g., ovulation and oviposition) (37, 41). However, our understanding of the molecular and physiological adaptations that underpin postmating female responses is taxonomically limited, impeding our understanding of basic insect physiology and sexual selection and development of tools to control insect reproduction (40).

The unusual reproductive system and novel paragenital female reproductive organs of bedbugs provides a unique opportunity to study the evolution of reproductive anatomy and postmating physiology. Bedbugs obligately mate by traumatic insemination, where the male pierces the female abdominal cuticle with modified external genitalia (called a paramere), transferring the ejaculate into the female body (50, 51). Intriguingly, female bedbugs have evolved a novel reproductive organ called the spermalege hypothesised to mitigate the costs of infection and wounding associated with mating (52, 53). The spermalege consists of several tissues and fulfils the copulatory and immune functions associated with the lower reproductive tract but is developmentally and anatomically separate (Fig. 1). In the common bedbug (*Cimex lectularius*), the mesodermally derived mesospermalege is the site of insemination, ejaculate × female interactions and reproductive immune activity (54–56). After insemination, sperm remain in the mesospermalege for several hours before crossing the tissue membrane, traversing the haemolymph, and entering into the female reproductive tract 3-6 hours after mating (51). Thus the bedbug female reproductive organs represent a system without the shared ontogeny and close localisation of female reproductive tract tissues that may otherwise constrain divergence in gene expression evolution (12).

**Figure 1.**
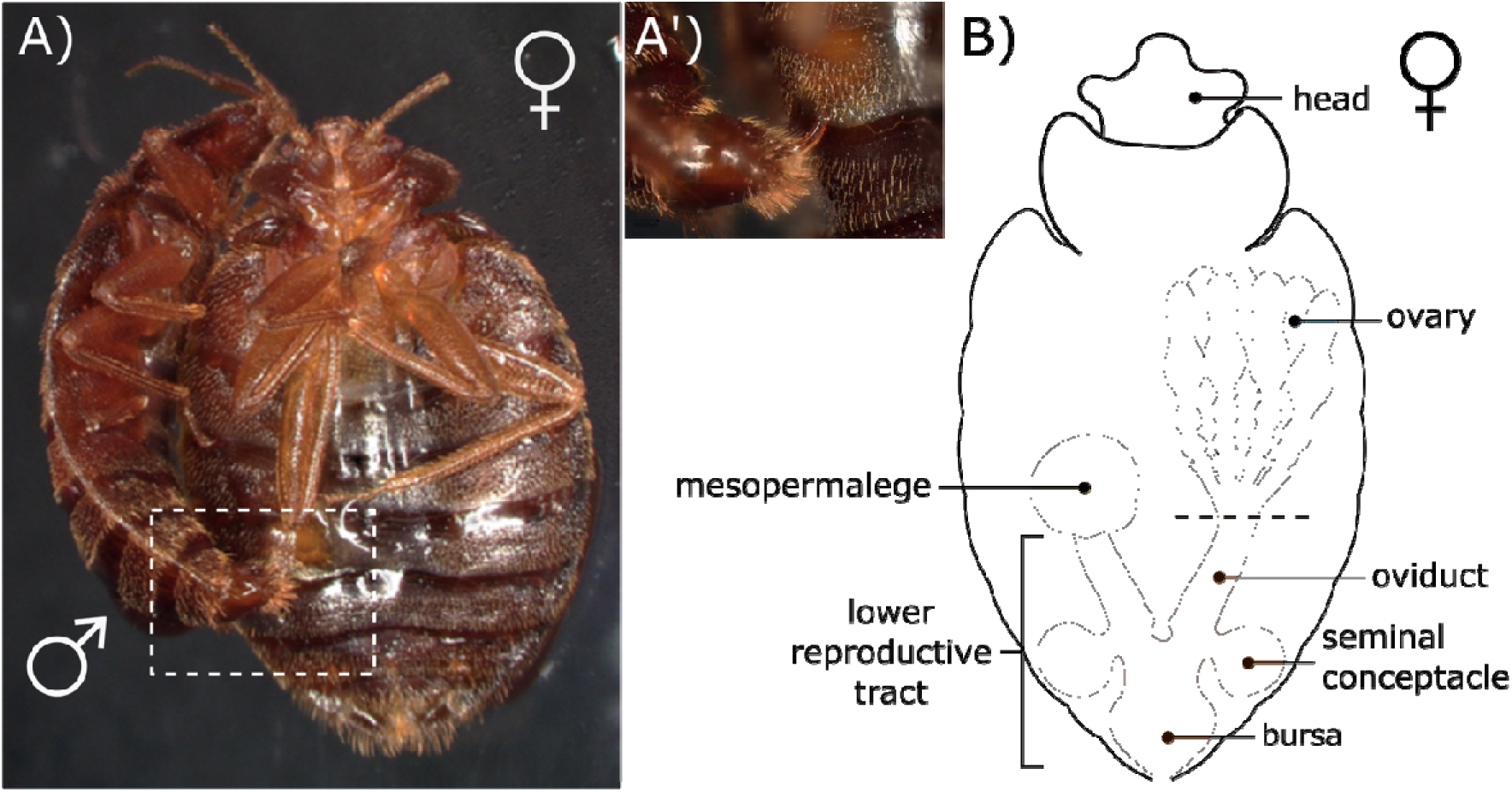
Traumatic insemination in the bedbug Cimex lectularius. **A)** Mating pair (ventral view) showing male (left) bending abdomen to pierce the female ectospermalege. **A’)** Close-up of paramere piercing the ectospermalege. **B)** Schematic female reproductive system (ventral view) adapted from (51). The lower reproductive tract (orange) comprises the bursa, seminal conceptacles and oviducts. Ovaries (blue) were separated during dissection at the dashed line. The mesospermalege (yellow) is fully distinct and not connected to the lower reproductive tract (right ovary omitted for clarity). Heads (red) were separated from the rest of the body.

In this study, we investigate how the evolution of a novel organ has shaped the function and postmating transcriptional response in the female reproductive system of the bedbug. We show that different aspects of the typical insect female postmating responses are divided across the mesospermalege and lower reproductive tract in the bedbug. The mesospermalege has acquired many of the functions associated with the reproductive tract as the site of insemination, including enrichment of genes involved in proteolysis and the immune response and expression of male seminal fluid genes. In parallel, the lower reproductive tract may have secondarily lost some of these features, instead being more specialised for interactions with sperm or facilitation of ovulation and oviposition. The postmating response in the mesospermalege and lower reproductive tract are mostly independent, with the postmating transcriptional response in the lower reproductive tract delayed until several hours after mating, coinciding with receipt of ejaculate (51). The divergent functions of differentially expressed genes further indicate a decoupling of reproductive tract functions between the two organs. Our study in a true bug (Hemiptera) with unique postmating physiology represents a notable expansion in our understanding of insect reproduction and the transcriptional evolution of a novel reproductive organ.

## METHODS

### Experimental animals

Adult bedbugs (*Cimex lectularius*) were collected from an urban infestation in London (UK) in 2006 and maintained as a large, outbred population with overlapping generations at TU Dresden. After eclosion, adults were kept in single-sex groups of seventy in 60 ml vials containing filter paper for two weeks. Bedbugs were kept at 26°C and 70% relative humidity on a 12:12 hour light-dark cycle and fed on human blood weekly to satiation following long-term experimental protocols (53).

Matings were conducted by pairing two-week old unmated females and males (one to two days after feeding) individually in 5.5 cm petri dishes lined with filter paper. We snap-froze unmated females and males in liquid nitrogen and mated females within seconds after mating (0h), 1h, 3h, 6h, or 24h after mating.

### Sperm transit through the female reproductive system

We dissected females from each timepoint (n = 5), imaged mesospermalege and reproductive tract, and scored for presence or absence of sperm. We also counted the numbers of sperm (stained with DAPI-PTW mix) in the mesospermalege contents over the time course after mating (n = 20). Measurements were performed blind to mating status and time postmating (see supplementary material).

### RNA extraction and sequencing

Bedbugs stored frozen at −80°C were thawed on ice and dissected and washed in ice-cold 1XPBS. We dissected heads from the body using fine surgical scissors. For males, we dissected the testes, separating and discarding the closely associated mycetomes. For female reproductive tissues, we isolated the female reproductive tract and mesospermalege into separate drops of PBS and removed surrounding fat body and any ejaculate mass present. The ovaries were then separated from the lower female reproductive tract and placed into a separate drop of PBS. Pooled tissues from ten individuals were stored in 50 µl RNAlater™ at 4°C until RNA extraction. We extracted total RNA with the Lucigen MasterPure™ Complete DNA and RNA Purification Kit with TURBO™ DNase treatment following standard protocols. Isolated RNA was resuspended in 30 µL HyClone™ HyPure water and stored at −80°C before sequencing. Library preparation and next-generation sequencing were performed at Azenta Life Sciences (Leipzig, Germany) using polyA-mRNA selection and paired-end sequencing of 150 bp reads on the Illumina NovaSeq X Plus sequencing platform (see supplementary material).

### Filtering, alignment, and quantification

We used the Nextflow nf-core/rnaseq pipeline (57, 58) to perform quality control, trimming, alignment, and quantification of sequenced reads. Reads were aligned to the *Cimex lectularius* reference genome (Clec_2.1, (59)) with STAR, (60) and quantified with Salmon (61). Read counts were imported into *R* using *tximport* (62). We removed genes with fewer than five counts per million reads (cpm) in fewer than three replicates using *edgeR* (63). To determine the chromosomal distribution of genes, we used BLASTn (77) to align coding sequences from the *C. lectularius* reference genome to the recently published chromosome-level *C. lectularius* genome assembly (78). For details see Supplementary Material.

### Tissue specificity and differential gene expression analysis

We calculated tissue-specific expression (*τ*) (65) using transcripts per million reads (TPMs) from unmated ovaries, lower female reproductive tracts and mesospermaleges, with unmated female and male heads and testes as outgroups. We excluded genes with TPM < 1 averaged across replicates (combined average for female and male heads) and classified genes as tissue-specific where *τ* ≥ 0.85 (Fig. S1). To analyse postmating transcriptional responses, we performed pairwise differential gene expression analysis comparing each postmating timepoint to unmated samples for each tissue separately using *DESeq2* (66) and log-fold-change shrinkage using *apeglm* (67). Genes were classified as differentially expressed based on a log2-fold-chanage > |1.5| and adjusted *p*-value < 0.05.

### Signal peptide identification and gene ontology enrichment analysis

We downloaded protein sequences for the entire bedbug proteome from uniprot.org and identified proteins containing predicted signal peptide sequences using *SignalP 6.0* (69) and *Phobius* (70), and combined the resulting lists. We matched UniProt IDs to NCBI IDs using DAVID (71, 72). We performed gene ontology (GO) enrichment analysis with *topGO* (68) using Fisher tests/weight01, nodeSize = 15. As background, we used the entire *C. lectularius* gene list for tissue-specific expression analysis (n = 20,808) or the genes expressed in that tissue (TPM ≥ 1) for the postmating response analysis (mesospermalege, n = 7,727; lower reproductive tract, n = 7,918). We report enriched terms with *p*-values < 0.05.

### Ortholog identification and evolutionary dynamics

We identified orthologs between *Cimex lectularius* and *Drosophila melanogaster* proteins downloaded from uniprot.org with *OrthoFinder* using default settings (73, 74). We calculated pairwise non-synonymous (dN) to synonymous (dS) nucleotide substitution rates between *C. lectularius* and the tropical bedbug (*C. hemipterus*) using PAML (75) using CDS for *C. lectularius* from NCBI (59) and the latest genome and GFF3 annotation for *C. hemipterus* (76). We used *ChatGPT* (GPT-4) to help generate *Python* scripts for these analyses. All scripts and results were inspected manually.

### Predicting protein-protein interactions using AlphaFold3

We used AlphaFold3 (79) to model protein-protein interactions between female mesospermalege-specific genes (n = 179) or lower reproductive tract genes (n = 112) encoding proteins containing a signal peptide sequence with putative *C. lectularius* male seminal fluid proteins (n = 249) (Garlovsky *et al.* in prep.). We used Claude (Anthropic, Sonnet 4.5) to optimise the AlphaFold3 pipeline for our HPC infrastructure. We subset proteins showing evidence of confident protein-protein complexes with interface predicted template modelling (ipTM) scores ≥ 0.8. For details see Supplementary Material.

### Statistical analysis

All statistical analysis was performed in R v4.5.0 (64). Full details of the statistical analyses see supplementary material and online code repository (https://github.com/MartinGarlovsky/Bedbug_RNAseq).

## RESULTS AND DISCUSSION

### Characterising mesospermalege and lower reproductive tract genes

The unique transcriptional characteristics of the bed bug mesospermalege, lower reproductive tracts (including the bursa, seminal conceptacles, and oviducts), heads and ovaries were identified by a principal component analysis (PCA) (Fig. 2A; Fig. S2A-C). The first PC (54% variance explained) separated the ovaries from other tissues, while PC2 (25.6% variance explained) clearly separated the novel mesospermalege organ from other reproductive tissues.

**Figure 2.**
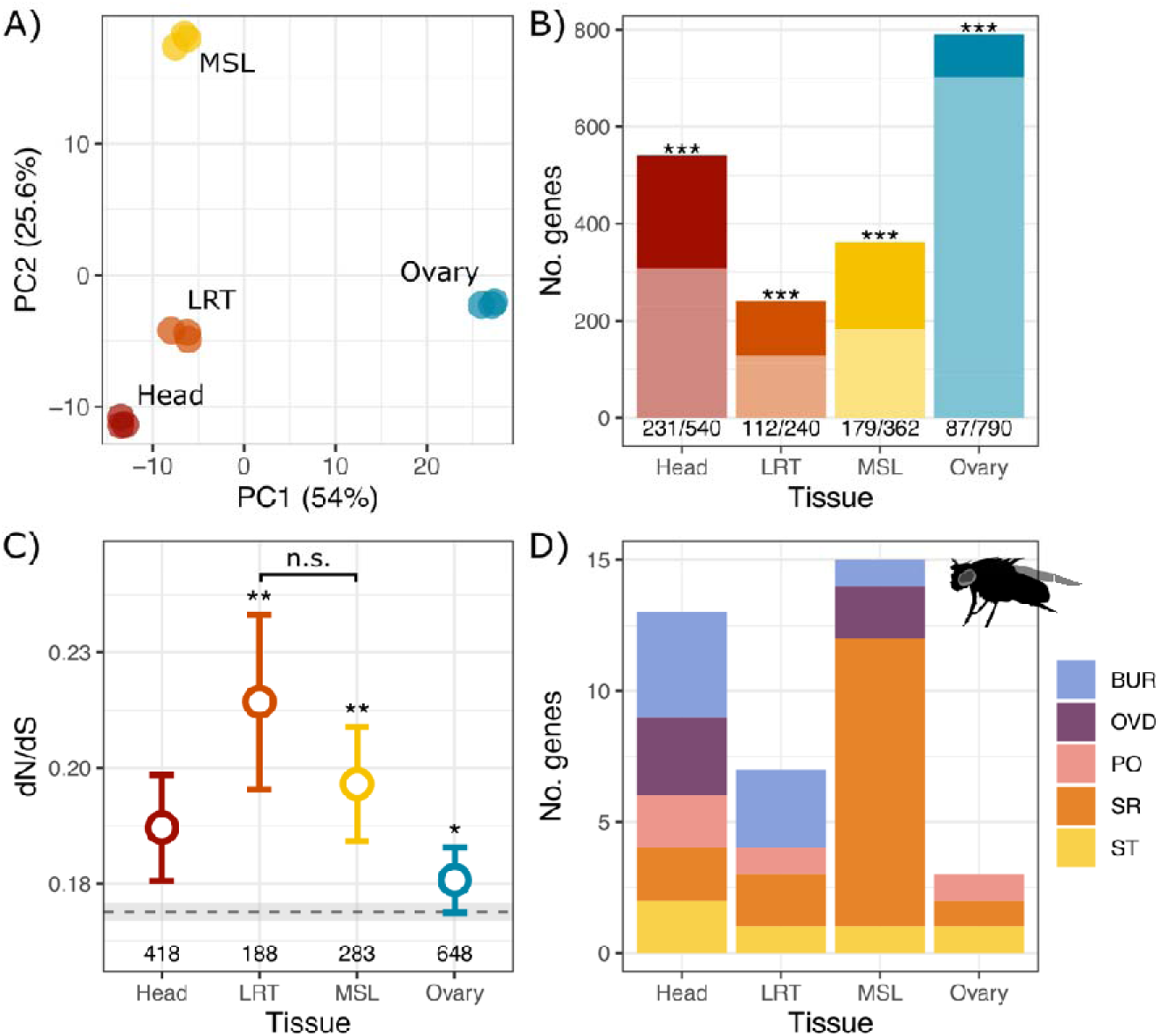
Characterising gene expression in the bedbug reproductive system. **A)** Principal component analysis of gene expression log2(normalised counts) for the 500 most variable genes from female heads, ovaries, mesospermaleges and lower reproductive tracts. **B)** Numbers of tissue-specific genes (τ ≥ 0.85) in each of the tissues. Darker shading shows number of genes encoding proteins containing a signal peptide sequence. Asterisks represent tissues with significantly different observed number of genes with a signal peptide sequence test) compared to the expected number in the genome. **C)** Molecular evolutionary rates (pairwise dN/dS, mean ± standard error) for genes with tissue-specific expression. Numbers below points indicate numbers of genes in each category. Asterisks represent results from Mann-Whitney U tests comparing each gene set to the genome average shown by dashed line (mean dN/dS = 0.17 ± 0.002, n = 9793). **D)** Numbers of tissue specific genes with D. melanogaster female reproductive tissue orthologs from (12): BUR, bursa; OVD, oviduct; PO, parovaria; SR, seminal receptacle; ST, spermatheca. n.s., not significant; *, p < 0.05; **, p < 0.01; ***, p < 0.001 (corrected for multiple testing using the Benjamini–Hochberg method).

Genes with specific expression are indicative of functional specificity in a tissue. Although all tissues expressed similar numbers of genes, the mesospermalege and lower reproductive tract expressed fewer tissue specific genes (*τ* ≥ 0.85) than the head or ovary (Fig. 2B, Fig. S1). However, this disparity can be explained by similarities between the two reproductive tissues as recalculating tissue-specificity for a combined female reproductive tissue category (average of mesospermalege and lower reproductive trat) showed similar numbers of tissue-specific genes compared to head or ovary (Fig. S3). There was a significant enrichment for a signal peptide sequence in mesospermalege genes (Χ^2^ = 94.3, df = 1, *p* < 0.001) and lower reproductive tract genes (Χ^2^ = 143.0, df = 1, *p* < 0.001) as well as heads (Χ^2^ = 309.30, df = 1, *p* < 0.001), indicative of proteins secreted into the tissue lumen (Fig. 2B). Ovary specific genes showed an under-enrichment of signal peptide sequences (Χ^2^ = 36.48, df = 1, *p* < 0.001). Neither the mesospermalege nor the lower reproductive tract showed enrichment of genes on either of the two *C. lectularius* X chromosomes (Fig. S4) (80), consistent with patterns in *Drosophila* (12, 39). Finally, both mesospermalege genes and lower reproductive tract genes were evolving faster than the genome average (Mann-Whitney *U*-test, both *p* < 0.001) but did not differ from each other (*p* = 0.798). Ovary-specific genes were also evolving rapidly (*p* = 0.01), whereas head-specific genes (*p* = 0.601) were evolving at a similar rate to the rest of the genome (Fig. 2C). The fast-paced molecular evolution of female reproductive genes that potentially interact with the male ejaculate is characteristic of sexually antagonistic selection (e.g., (81) but see (82)). Together, these results suggest the mesospermalege and lower reproductive tract experience similar evolutionary pressures.

#### Mesospermalege-specific genes

Consistent with mesospermalege functions in mitigating the costs of mating in general, and traumatic insemination in particular, mesospermalege-specific genes showed enrichment of immune response functions, including defense response, endo- and exo-cytosis, response to external stimulus and lysosome organelles (Table S1). These mesospermalege-specific genes included orthologs of genes expressed in the *D. melanogaster* female reproductive tract (12) involved in wound healing and ovulation (*matrix metalloproteinase 2, prophenoloxidase 1, hemolectin*), the innate immune response (*Tollo*; (83), lysosomal protein degradation *Cathepsin B* (*CtsB*), and haemocyte maturation (*Rho-like*) (Fig. S5).

Mesospermalege-specific genes also showed enrichment of many GO terms associated with female reproductive tract tissues in other species (40), consistent with the mesospermalege being the site of insemination. For instance, we found enrichment of GO terms including proteolysis, negative regulation of peptidase activity, and G protein-coupled receptor activity (Table S1). Proteases are frequently found in the reproductive tract and associated with breakdown of the ejaculate that facilitates sperm movement (e.g., *Drosophila* (84), cabbage white butterflies (25) and mice (85)). Interestingly, an ortholog of the *sex peptide receptor* (*SPR*) gene, a G protein-coupled receptor that induces the postmating switch in *D. melanogaster* (86), showed highest expression in the mesospermalege rather than the lower reproductive tract (Fig. S5).

Overall, mesospermalege-specific genes included more orthologs of *D. melanogaster* female reproductive tissue specific genes than the lower reproductive tract (12) (Fig. 2D). Among the orthologs of *D. melanogaster* genes, we identified an odorant binding protein (Odp57c) involved in postmating responses (87) and genes involved in synaptic/signal transmission (e.g., *CaMKII*, *Hykk*, *atilla*, *pigs*, *tnc*, *NPFR*, *nAChR*β*3*, *VGlut2*) (Table S2; Fig. S5). This is consistent with the mesospermalege having evolved an expression profile like the lower female reproductive tract of other species despite its independent developmental origins. This pattern also provides further evidence that the functions involved in response to intromission and ejaculate processing drive convergent evolution of reproductive tract gene expression across diverse species.

#### Lower reproductive tract-specific genes

Showed enrichment of GO terms related to transport and localisation, metabolic and catalytic processes, regulation and signalling (Table S2). We identified several orthologs of *D. melanogaster* genes involved in reproduction, including *hadley*, a member of the *sex peptide* network involved in negative regulation of female receptivity postmating (88); *Neprilysin 1*, expressed in *Drosophila* nervous system and spermathecae, essential for female fertility (89); and genes involved in metabolism (*Aldose reductase 7*, *Curly Su, Pxn*), signalling (*mind the gap*, *TpnC73F*, *egr*, *trol*) and transport (*Picot*, *Best2*) (Table S2). Although the lower reproductive tract shows similar enrichments to some properties of the female reproductive tract in other species (e.g., metabolism) common enrichments of genes with immune response and gustatory receptors are absent. Instead, enriched functions in the lower reproductive tract may more directly relate to lower reproductive tract functions in interactions with sperm or oviposition and ovulation.

#### Seminal fluid protein expression in the female reproductive organs

Evidence of postmating cooperation between the sexes has been suggested because of the shared expression of male seminal fluid proteins in the female reproductive tract in *Drosophila* (12, 39, 90). In unmated females we found expression (TPM ≥ 1) of most (87.6%; 218/249) putative bedbug seminal fluid proteins (Garlovsky et al. in prep.). These seminal fluid proteins tended to have highest expression in the mesospermalege, including 19 out of 46 seminal fluid proteins showing tissue-specific expression (Fig. 3A). This pattern further indicates that the mesospermalege has taken on the functions associated with ejaculate processing in the bedbug. Moreover, identification of seminal fluid genes in the bedbug female reproductive system demonstrates that this pattern is not only found in *Drosophila* and may represent a more common pattern of intersexual dynamics in reproductive system gene expression evolution.

**Figure 3.**
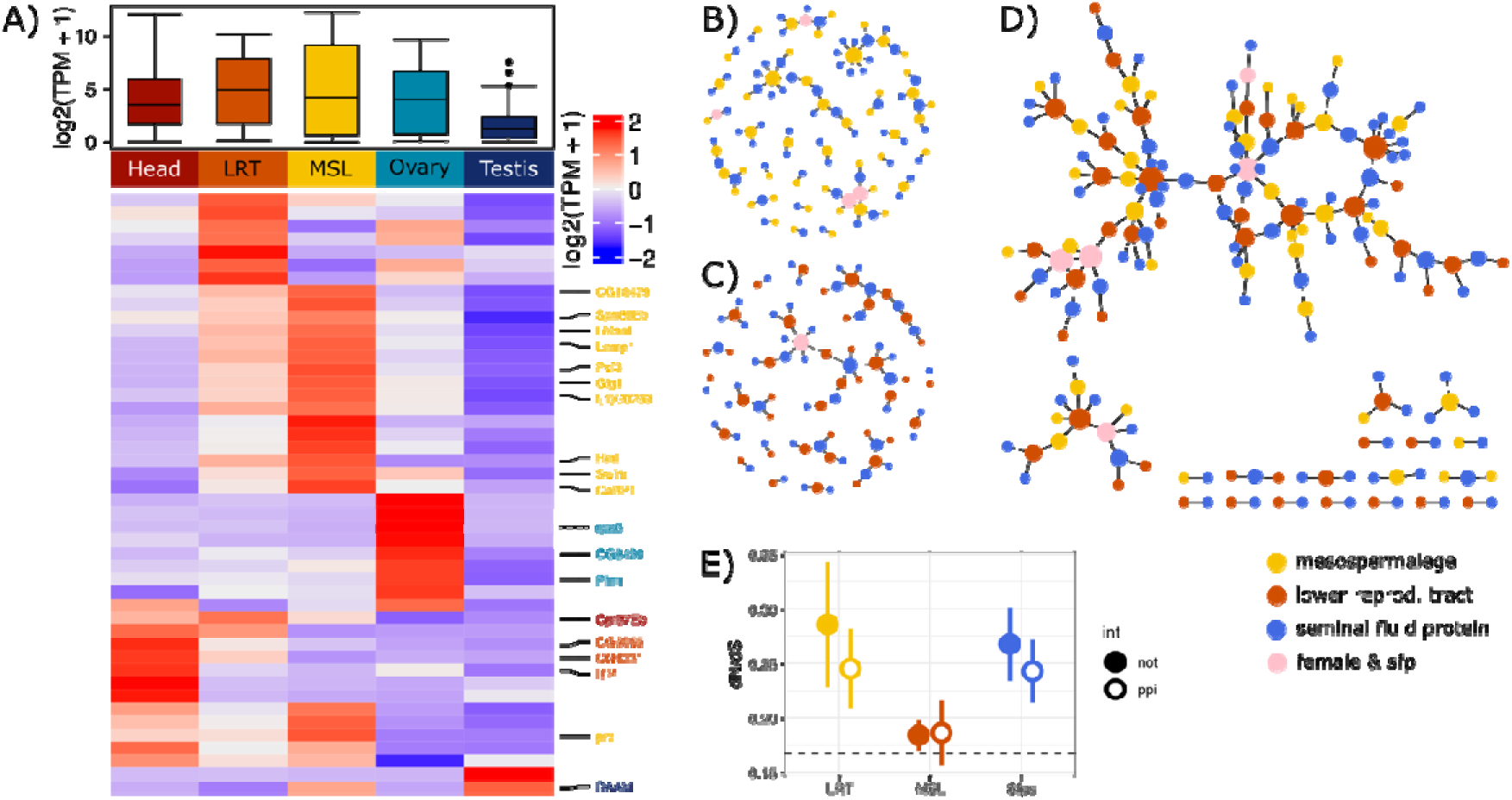
**A)** Expression of putative male seminal fluid proteins in female reproductive tissues (n = 46). Boxplots (top) show log2(TPM + 1) in each tissue. Heatmap (bottom) shows normalised expression log2(TPM + 1) across each tissue. D. melanogaster orthologs are labelled on right hand side and coloured based on tissue-specific expression. MSL: mesospermalege, LRT: lower reproductive tract. **B)** Network diagram for mesospermalege (yellow) and seminal fluid protein (blue) interactions (ipTM ≥ 0.8). **C)** Network diagram for lower reproductive tract (red) and seminal fluid protein (blue) interactions (ipTM ≥ 0.8). Some proteins were identified as both a mesospermalege/lower reproductive tract protein and a male seminal fluid protein (pink). **D)** Network diagram combining mesospermalege- and lower reproductive tract-interactions with seminal fluid proteins (combining B and C). **E)** Molecular evolutionary rates (pairwise dN/dS, mean ± standard error) for genes with (open) or without (filled) evidence of interaction.

### Ejaculate-female interactions

Given the abundance of female reproductive genes containing a signal peptide sequence (Fig. 2B), we hypothesised that these putatively secreted proteins interact with the male seminal fluid proteome. Using *in-silico* predications we found one fifth (20.9%; 52/249) of putatively secreted mesospermalege proteins showed evidence of a protein-protein interaction (ipTM ≥ 0.8) with at least one seminal fluid protein (Fig. 3B). Contrary to our hypothesis that a majority of ejaculate processing occurs in the mesospermalege, a greater proportion of lower reproductive tract proteins showed evidence of an interaction (36.6%; 41/112; Fisher’s exact test, *p* = 0.020). Interacting female proteins included several orthologs of *Drosophila* proteins with roles in postmating interactions (see above), including genes in the Sex Peptide network (Table 1). Overall, these interacting proteins were not evolving faster than the rest of the secretome (Fig. 3E). This suggests specific female-male protein interactions are not responsible for the elevated rate of molecular evolution found in female reproductive genes.

**Table 1.**
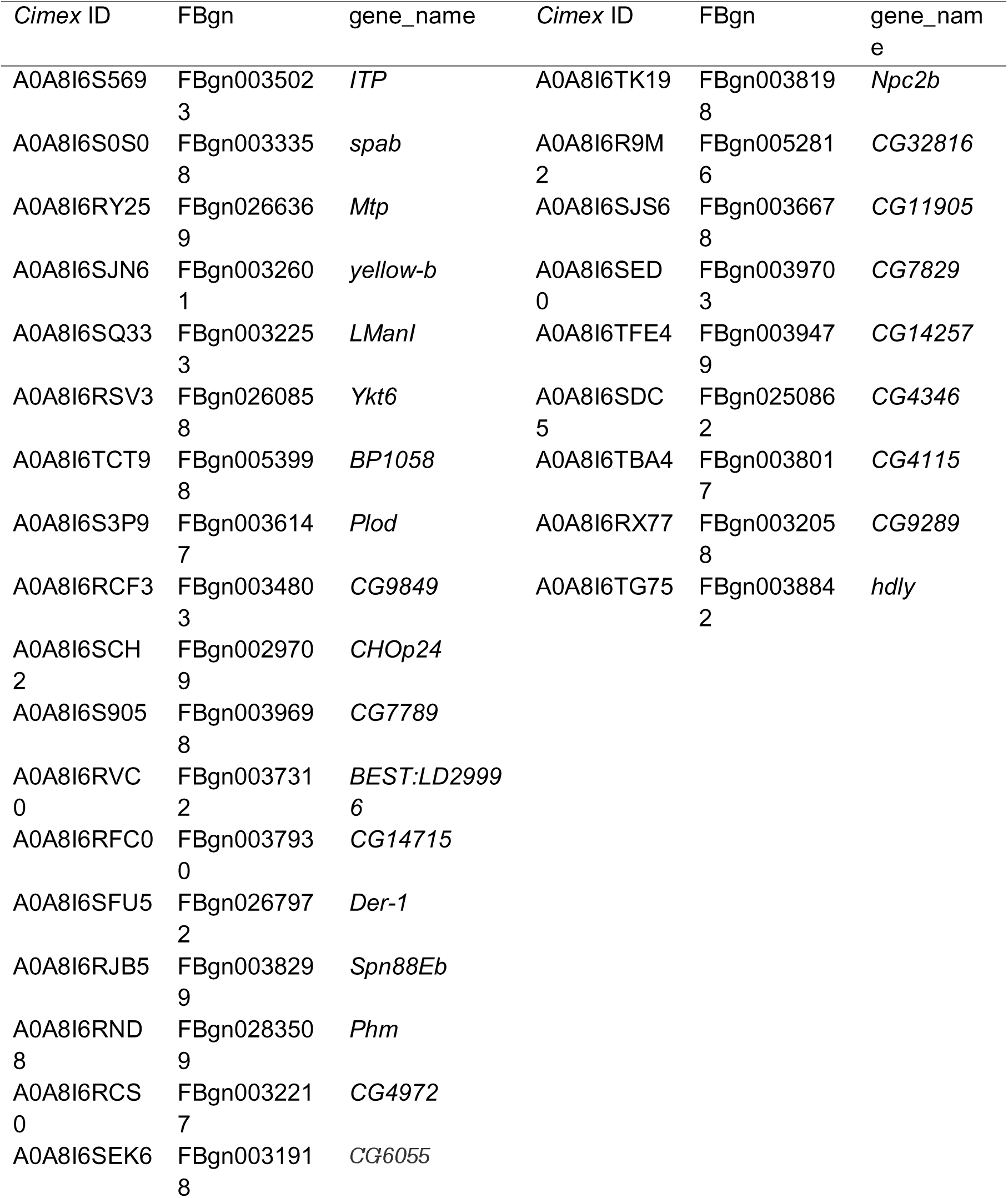
Female reproductive proteins that interact with the bedbug male seminal fluid proteome with Drosophila melanogaster orthologs.

### Postmating transcriptional responses in the mesospermalege and lower reproductive tract

We next compared the postmating transcriptional response in the mesospermalege and lower reproductive tract over the first 24h after mating. Immediately after mating the mesospermalege is filled with the sperm mass while there are no visual changes in the lower reproductive tract (Fig. 4A). Sperm numbers rapidly decreased after mating (GLM, F_1,99_ = 144.77, *p* < 0.001) falling significantly to 50% of starting concentration 1h after mating (Welch two-sample t-test, *t* = 3.38, df = 19.55, *p* = 0.003) as sperm begin to exit *enroute* towards the reproductive tract (Fig. 4B). The reduction in both sperm numbers and variance (F-test, F_19,19_ = 13.22, *p* < 0.001) may result from some sperm in some individuals immediately leaving the mesospermalege or being digested by the abundant haemocytes in the mesospermalege. Sperm begin to arrive in the seminal conceptacles as early as 1h after mating (Fig. 4C). By 6h after mating sperm numbers in the mesospermalege reduced by another 50%, with no or very few sperm present by 24h after mating, while the lower reproductive tract becomes full of sperm (Fig. 4B&C).

**Figure 4.**
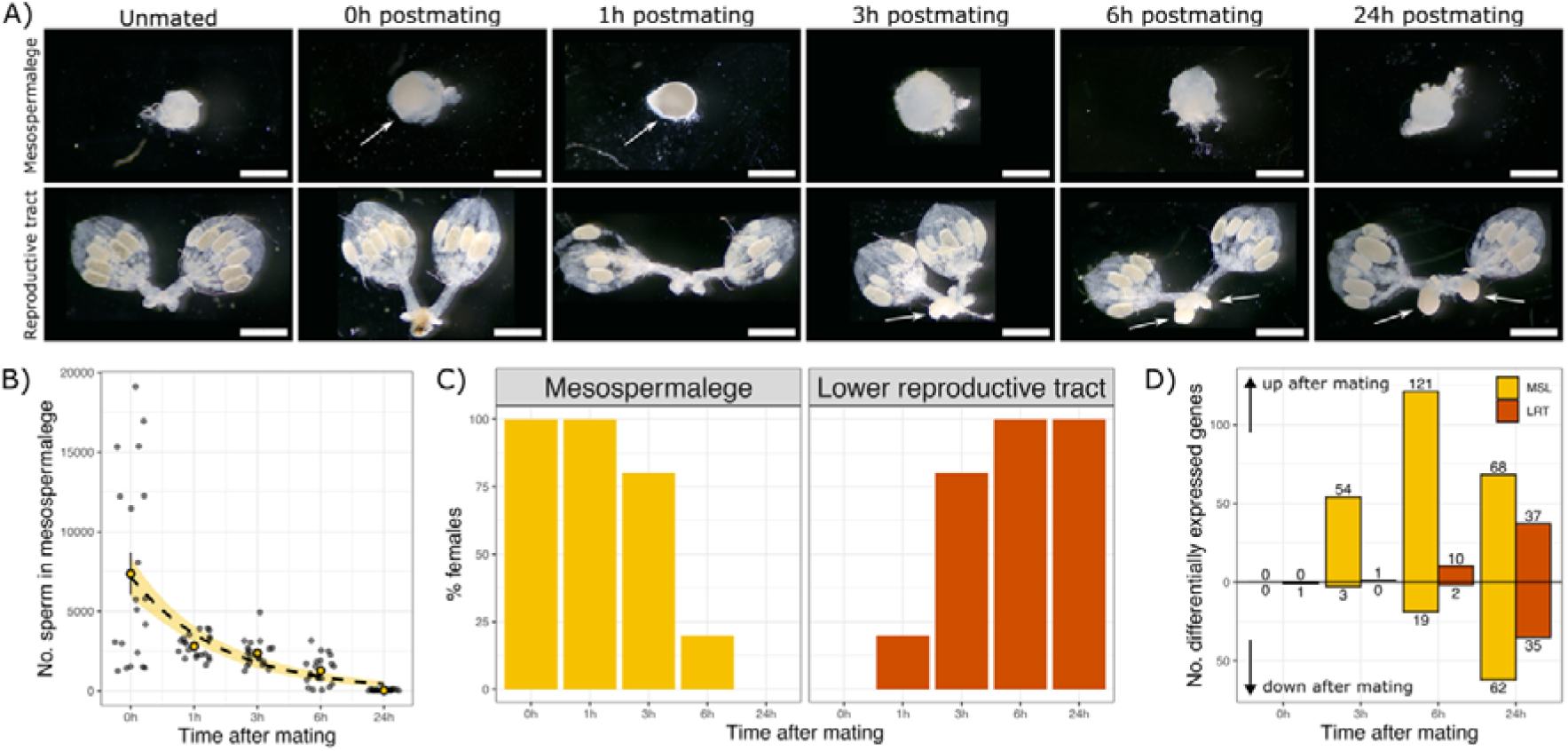
Sperm transit through the female reproductive system. **A)** Representative images of mesospermalege (top) and reproductive tract (bottom) at each timepoint. Sperm mass indicated by arrows (Scale bar, 1mm). **B)** Numbers of sperm counted in the mesospermalege after mating. Large points are means ± standard error and smaller points are individual females. The dashed line represents the estimate (± 95% confidence interval) from a GLM. **C)** Percentage of females with sperm observed (present/absent) in mesospermalege (left; yellow) and lower reproductive tract (right; orange). **D)** Number of differentially expressed genes up- or down-regulated in the mesospermalege (MSL; yellow) and lower reproductive tract (LRT; orange) compared to unmated samples. Numbers above/below bars indicate the number of genes up- and down-regulated, respectively.

Corresponding to their different temporal interactions with the ejaculate, the mesospermalege had an earlier postmating transcriptional response than the lower reproductive tract (Fig. 4D; Fig. S6&7). The two tissues also exhibited marked differences in their patterns of increased and decreased expression with initial changes in gene expression being primarily upregulation in the mesospermalege and a mix of up and downregulation in the lower reproductive tract.

The majority of the postmating transcriptional response in the mesospermalege was unique, with only 21 genes (8.4%) that were also differentially expressed in the lower female reproductive tract (Fig. S8A). These genes showed no correlation in expression at earlier timepoints, indicating the organs are responding independently shortly after mating. By 6h postmating and later, expression of these 21 genes was positively correlated (Fig. S8B). On the other hand, these 21 genes make up over a quarter (26.9%) of differentially expressed genes in the lower reproductive tract, suggesting a core of common postmating functions between the tissues.

#### Postmating response in the mesospermalege

The mesospermalege only began to show a postmating transcriptional response 3h after mating (Fig. 4D & Fig. S7). Lack of differential expression in the mesospermalege immediately after mating (0h) suggests that perception of a male or mating do not initiate an immediate postmating female response (49). Most differentially expressed genes at 3h and 6h postmating were upregulated, indicating receipt of the ejaculate “switches on” transcription in the mesospermalege. The peak in differential expression occurred at 6h postmating, similar to the timing in *D. melanogaster* (12, 38). By 24h after mating, the number of downregulated genes increased, suggesting repression of transcriptional activity once sperm have left the mesospermalege (Fig. 4). Clustering of genes (k-means = 5) with similar postmating expression profiles identified groups of genes with either decrease (2 clusters) or increase (3 clusters) over the time course (Fig 5).

**Figure 5.**
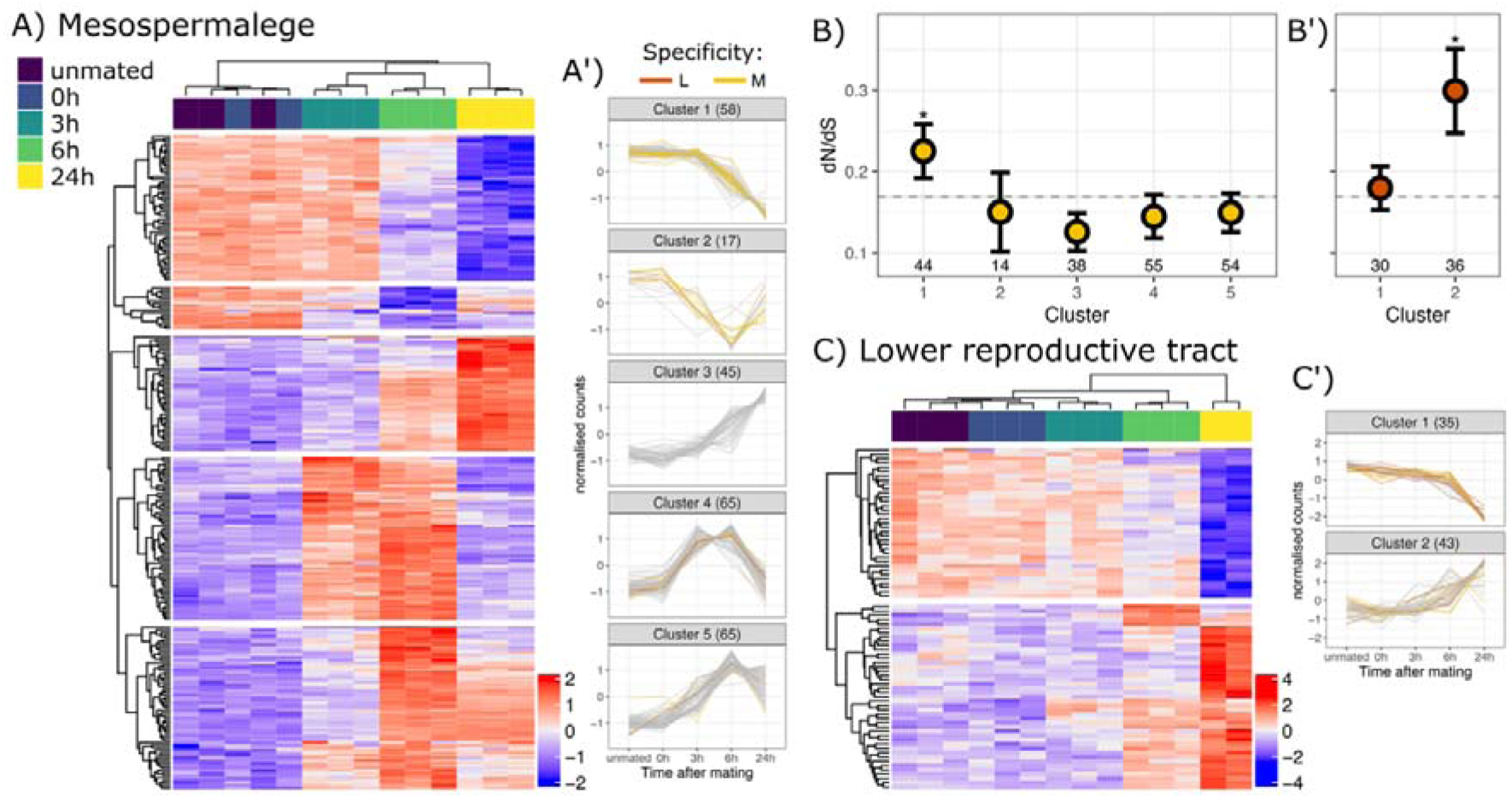
Postmating transcriptional response in the mesospermalege and lower reproductive tract. **A)** Heatmap of normalised expression log2(n + 1) of differentially expressed genes in the mesospermalege (n = 251). **A’)** Mean normalised expression for each gene (lines) in each K-means cluster (k = 5) coloured by tissue-specific expression (L = lower reproductive tracts, M = mesospermalege). **B)** Molecular evolutionary rates (mean dN/dS ± standard error) of genes showing differential expression after mating in the mesospermalege (k = 5) and **B’)** lower reproductive tract (k = 2). Numbers of genes are indicated below points. Asterisks represent results from Mann-Whitney U tests comparing each gene set to the genome average shown by dashed line (mean dN/dS = 0.17 ± 0.002, n = 9793). **C)** Heatmap of normalised expression log2(n + 1) of differentially expressed genes in the lower reproductive tract (n = 78). **C’)** Mean normalised expression for each gene (lines) in each K-means cluster (k = 2). Colours in C and C’ as in A and A’.

Genes in clusters with increased expression likely contribute to postmating responses in the female. The earliest increases in postmating expression were transient (Cluster 4) and enriched for genes involved in biosynthesis and catabolism with transcriptional regulators and membrane transporters. Genes with a later stable or transient increase in expression (Cluster 5) showed enrichment of genes including methyltransferases, RNA-modifying enzymes, and energy-dependent regulators coordinating gene expression and developmental signalling. Finally, genes with consistent increase in expression after mating (Cluster 3) were enriched for RNA processing and translation (Table S3). As most of these gene expression changes occur after the ejaculate exited the mesospermalege (Fig 4) these changes likely reflect a metabolic and functional restoration of the tissue.

Genes downregulated after mating showed an enrichment of mesospermalege-specific genes (Cluster1: 44.8%, 26/58; Cluster 2: 35.3%, 6/16) (Fig. 5A’). Genes in Cluster 1, which decrease in expression in all postmating timepoints were enriched for GO terms relating to signal transduction and synaptic communication, receptor and channel activity, transporter activity, catalytic and redox activity and molecule binding (Table S3). Notably, these genes showed elevated rates of molecular evolution (Mann-Whitney *U* test, *p* = 0.021; Fig. 5B), which along with their expression pattern supports their function in ejaculate processing and sexual conflict (91). Genes in Cluster 2, which decrease until 6h postmating and then return towards the unmated state, were involved in the regulation of signalling and cell cycle progression (Table S3).

Although greater postmating upregulation of gene expression over the course of mating has been the dominant pattern in insect studies, especially in *D. melanogaster* (12, 37–39, 92), there are notable exceptions that also show high levels of down regulation. For example, *Drosophila mojavensis* also showed more decreased expression at later postmating timepoints (93). Notably, a large insemination reaction forms in the female reproductive tract after mating in the *mojavensis* group which looks somewhat like the sperm mass in the mesospermalege immediately after mating (Garlovsky, pers. obs.). The *mojavensis* insemination reaction involves high protease expression and may result in a dominance of ejaculate processing functions in postmating expression patterns making it more similar to the mesospermalege (94, 95). Similarly, the initial function in the digestion of the hard outer casing of the spermatophore in the cabbage white butterfly coincides with a decrease in expression of genes including proteases likely involved in initial interactions with the ejaculate (25). The mesospermalege therefore shows a postmating transcriptional response characteristic of functions associated with response to mating and ejaculate processing. Namely, an earlier peak in expression differences and a decrease in genes that are necessary for ejaculate-female interactions but may have broad digestive functions that may be detrimental for tissue integrity if elevated long term (25, 26).

#### Postmating response in the lower reproductive tract

In the lower reproductive tract differential gene expression started later. Substantive numbers of differentially expressed genes were only identified 24h after mating (Fig. 4B & 4C & Fig. S7). These genes fit two clusters (k = 2); those that are increasing or decreasing in expression. Genes downregulated after mating (Cluster 1; Fig. 5C’) were enriched for metabolic and biosynthetic processes, secretion, defense response, neuron differentiation; extracellular region, membrane, and anchoring junction; lipase activity and receptivity and transporter activity (Table S4). Genes upregulated after mating (Cluster 2) were enriched for negative regulation of signal transduction, response to oxidative stress, defense response; extracellular region; lipid binding (Table S4). Contrary to the pattern seen in the mesospermalege, lower reproductive tract genes that increased in expression evolved faster than the genome average (*p* = 0.035; Fig. 5B) indicative of rapidly evolving reproductive genes involved in interactions with the ejaculate. However, the delayed pattern of differential expression in the lower reproductive tract will require investigating timepoints beyond the first 24 hours after mating to evaluate the full trajectory of gene expression changes.

This early snapshot of the bedbug lower reproductive tract nevertheless reveals a markedly different pattern from *D. melanogaster* which shows a peak in differential expression at 6h postmating (12, 38). Evolvability of postmating gene expression changes to correspond to different functional timings is supported by patterns in other species. For example, in the *Drosophila virilis* species complex the peak in postmating differences occurs earlier (3h versus 6h), which may relate to differences in the timing of reproductive events (39). In contrast, major postmating expression changes are not observed in the lower reproductive tract of cabbage white butterflies until 72h postmating, corresponding to the delayed opening of the spermatophore (25). A later peak in postmating response (24h) is also found in mosquitos which may correspond to the longer time required for females to produce and lay their first egg batch (37). Thus, the postmating transcriptional response in the lower reproductive tract is consistent with evolutionary shifts in the timing of differential expression to coincide with the functional requirements of sperm entry into the reproductive tract (51).

## CONCLUSION

The specialisation of reproductive functions between the bedbug lower reproductive tract and the mesospermalege are evident in their transcriptional profiles. Higher expression of immune genes and proteases in the mesospermalege is consistent with its replacement of the lower reproductive tract in immune response following insemination and initial ejaculate processing. The rapid evolution of tissue specific genes, as well as specific expression of and interaction with seminal fluid protein genes, are indicative of distinct interactions between each tissue and the ejaculate. The specific postmating functions of the mesospermalege are further supported by an early peak in differential expression and a larger proportion of rapidly evolving tissue-specific genes that decrease in expression, consistent with patterns found in species with extensive ejaculate processing (25, 37). Gene expression in the lower reproductive tract, in turn, becomes more specific to sperm storage as well as processes of ovulation and oviposition. This can be seen in the delayed initiation of changes in postmating expression and rapid evolution of genes that increase postmating. The distribution of postmating ejaculate-female and reproductive functions across two tissues provides unique insights into the evolution of gene expression in novel tissues and postmating expression dynamics. We further identify broader relationships in the co-expression of seminal fluid genes in the female reproductive system and the relationship between postmating responses and the reproductive biology of the organism. Future research across species can take the patterns we have shown in the bedbug into consideration when evaluating what functions might contribute to postmating expression profiles. Further investigation, particularly with single cell technologies, will provide additional opportunities to explore the constraints on gene expression evolution of reproductive organs that are overcome by the evolutionary novelty of the mesospermalege in traumatic insemination.

## ACKNOWLEDGMENTS

We thank Alison E. Wright and Christoph-Rüdiger von Bredow for comments on an earlier version of the manuscript, Christin Froschauer for help in the lab, and Yasir H. Ahmed-Braimah for discussion about analysis. We are grateful for help and support from Yishai Gilron, Yan Ge and the ZIH team at TU Dresden and for computing time made available on the high-performance computer at the NHR Centre of TU Dresden.

## FUNDING

This project was funded by a TU Dresden Graduate Academy grant awarded to MDG, funded by the BMFTR and the Free State of Saxony under the Excellence Strategy of Federal Government and the Länder. CEMG was supported by a National Science Foundation Postdoctoral Research Fellowship in Biology (PRFB-2208973). The NHR Centre of TU Dresden is jointly supported by the Federal Ministry of Research, Technology and Space of Germany and the state governments participating in the NHR (www.nhr-verein.de/unsere-partner).

## Notes

https://doi.org/10.25532/OPARA-1133

### Competing Interest Statement

The authors have declared no competing interest.

