## Supplementary Materials for "Novel female reproductive organ differentiates postmating transcriptional response to insemination versus arrival of sperm in bedbugs"

**FUNDING**

This project was funded by a TU Dresden Graduate Academy grant awarded to MDG, funded by the BMFTR and the Free State of Saxony under the Excellence Strategy of Federal Government and the Länder. CEMG was supported by a National Science Foundation Postdoctoral Research Fellowship in Biology (PRFB-2208973). The NHR Centre of TU Dresden is jointly supported by the Federal Ministry of Research, Technology and Space of Germany and the state governments participating in the NHR ([www.nhr-verein.de/unsere-partner](http://www.nhr-verein.de/unsere-partner)).

**DATA AVAILABILITY**

The processed data and full details of the analysis will be made available on the Open Access Repository of Saxon Universities (<https://opara.zih.tu-dresden.de/home>) at <https://doi.org/10.25532/OPARA-1133> and GitHub (https://github.com/MartinGarlovsky/Bedbug_RNAseq). Fastq files are available through the Sequence Read Archive (SRA) under project accession PRJNA1427960.

**SUPPLEMENTAL METHODS**

***Experimental animals***

Adult bedbugs (*Cimex lectularius*) were collected from an urban infestation in London, UK in 2006 and maintained as a large, outbred population with overlapping generations at TU Dresden. After eclosion, adults were kept in single-sex groups of seventy in 60 ml vials containing filter paper for two weeks. Bedbugs were kept at 26°C and 70% relative humidity on a 12:12 hour light-dark cycle and fed on human blood weekly to satiation following long-term experimental protocols (1).

Matings were conducted by pairing two-week old unmated females and males (one to two days after feeding) individually in 5.5 cm petri dishes lined with filter paper. We recorded the start and end of copulation (mean copulation duration ± standard error = 156.7 ± 3.2 s; range: 60-360 s). Females that mated for less than one minute or more than ten minutes were excluded. We snap-froze unmated females and males in liquid nitrogen and mated females within seconds after mating (0h), 1h, 3h, 6h, or 24h after mating. Frozen individuals were stored at -80ºC until dissection.

***Timing of sperm presence in female reproductive tissues***

We dissected five unmated females and mated females for each timepoint (n = 5) in 1XPBS using a stereo microscope (Stemi 305, Zeiss, Germany). First, we removed the complete female reproductive tract by gently pulling the last abdominal segment away from the abdomen. We cleared the reproductive tract of attached trachea and fat body and washed them twice in 30 µl drops of PBS. Then, we transferred the reproductive tract to a fresh 90-mm Petri dish into a 30µl drop of Leibovitz L-15 medium on a two-layer coverslip bridge and covered with a microscope slide. Images were captured with a digital camera (Axiocam 212 color, Zeiss, Germany) mounted on a stereo microscope. The mesospermalege was processed in the same way. We recorded images at 3000x magnification. Images were scored for the presence or absence of sperm in the mesospermalege or lower reproductive tract (seminal conceptacles, bursa, or oviduct). Counts were performed blind to treatment.

***Sperm counts in the mesospermalege***

We dissected females in a drop of 1XPBS. The mesospermalege was transferred to a second drop of PBS and ruptured to release the sperm mass. We transferred the sperm mass to an Eppendorf tube, diluted it in PBS and vortexed it. We diluted samples to different final volumes to aid sperm counting (final volume 1000µL for 0h, 1h, 3h and 6h; 500 µL for 24h). We then transferred 20µL of each solution to a multi-well slide and dried for one hour with three samples per female. Slides were moistened by addition- and immediate removal- of 20µL PBS, incubated in 15µL DAPI-PTW mix (1XPBS + 0.1% v/v Tween 20) for 15 minutes, washed twice with 20µL PBS for 15 minutes each, and mounted with Fluoromount G (Thermo). We photographed slides using a Leica DMi8 microscope in brightfield and with a DAPI filter, with 6-second exposure, gain 2 and 100% intensity. We counted the numbers of sperm at three predefined locations for each sample in ImageJ (2). Sperm counts were performed blind to sample condition, and a subset of images were counted twice to estimate observer repeatability (Pearson’s correlation = 0.994, confidence intervals [0.993, 0.995]). Total numbers of sperm in the mesospermalege were calculated with respect to the final dilution volume of the sperm mass and averaged across the three technical replicates. We analysed sperm counts in the mesospermalege with a quasi-Poisson generalised linear model using mean sperm counts per female with time after mating as the response.

***RNA extraction and sequencing***

We determined RNA concentrations with the Qubit™ RNA High Sensitivity Assay Kit with a Qubit® 3.0 Fluorometer. RNA quality was assessed at Azenta Life Sciences (Leipzig, Germany) with a fragment analyser using PROSize v3.0 (Agilent). The RNA Quality Number (RQN) calculation is unreliable in *C. lectularius* due to the denaturation of the 28S rRNA (3). Instead, we visually inspected the electropherograms to determine RNA quality and subsequently excluded two samples from further analysis (one 0h mesospermalege; one 24h lower reproductive tract). We sequenced female and male heads and gonads (testes and ovaries) separately from mesospermalege and lower reproductive tract samples. For heads and gonads, we sequenced a total of 644,426,052 reads with a mean quality score of 35.21. For postmating time-series samples of mesospermalege and lower reproductive tract tissues, we sequenced a total of 310,531,797 reads with a mean quality score of 38.54.

***Characterising gene expression patterns and signal sequence enrichment***

We performed principal component analysis for the 500 most variable genes in unmated tissues using log2(normalised counts) per gene from *DESeq2* to account for different library sizes (15). We compared the observed number of genes with a signal peptide sequence to the expected number in the genome in each tissue-specific gene sets using $X^{2}$ tests. To compare postmating timepoints we performed principal component analyses of the 500 most variable genes in each tissue using log2(normalised counts).

***Chromosomal distribution***

To determine the chromosomal distribution of genes, we extracted coding sequences from the GTF and genomic FASTA from the *C. lectularius* reference genome Clec_2.1 (4) using gffread v0.12.7 (5). We aligned sequences to the recently published chromosome-level *C. lectularius* genome assembly (6) using BLASTn (7) and constructed a nucleotide BLAST database from the reference genome using *makeblastdb* and aligned sequences using *blastn*. We chose the top-scoring alignment (highest bit score) for each gene to determine chromosomal location. We compared the observed number of genes on each chromosome to the expected number for each tissue-specific gene sets using $X^{2}$ tests.

***Evolutionary rates***

We calculated pairwise nonsynonymous (dN) and synonymous (dS) substitution rates (dN/dS) between *C. lectularius* and the tropical bedbug, *C. hemipterius* using PAML (8). We obtained coding sequences (CDS) for *C. lectularius* from NCBI (9). For *C. hemipterus* gene models were extracted using gffread using the latest genome and GFF3 annotation (10). We retrieved the longest isoform of each gene and performed reciprocal best BLAST using blastn (BLAST+ v.2.14.0) and retained only reciprocal best hits. Protein sequences were aligned using MUSCLE v5.1 (11) and codon-aware alignments constructed using PAL2NAL v14 (12) with default options. We estimated dN and dS using codeml (PAML v4.10.7) in pairwise mode (runmode = -2). The model used one ω ratio across all sites (model = 0, NSsites = 0), with estimated values for κ and ω (fix_kappa = 0, fix_omega = 0, cleandata = 1). Pairs with extreme or unreliable dS values (dS = 0 or >5) were excluded to reduce error from saturation. We used *ChatGPT* to help generate *Python* scripts for these analyses. All analyses were checked manually. We compared rates of molecular evolution between gene set and the rest of the genes in the genome using Mann-Whitney *U* tests.

***Predicting protein-protein interactions using AlphaFold3***

We downloaded the *C. lectularius* proteome from uniprot.org and removed signal peptide sequences identified by *Phobius* and *SignalP* 6.0 in *R* using the *Biostrings* package (18). We used *Claude* (Anthropic, Sonnet 4.5) to optimise the AlphaFold3 pipeline for our HPC infrastructure. All results were inspected manually. Less than 1% of protein pairs (mesospermalege-seminal fluid proteins: 54/44,571, 0.12%; lower reproductive tract-seminal fluid proteins: 38/27,888, 0.14%) did not fold due to large token size. We used *igraph* (19), *ggraph* (20), and *tidygraph* (21) to visualise protein networks for the subset of proteins with interface predicted template modelling (ipTM) values ≥ 0.8 which indicate confident model prediction of an interaction.

**SUPPLEMENTAL RESULTS**

Overall, we identified similar numbers of genes expressed (TPM ≥ 1) in each tissue (Fig. S1).

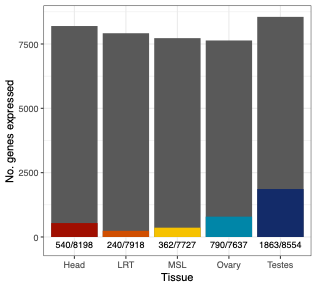

***Figure S1.*** *Number of genes expressed in each tissue with a minimum TPM > 1 are shown by the full height of each bar. The filled portion in colour shows the number of tissue-specific genes (τ ≥ 0.85) in each tissue.*

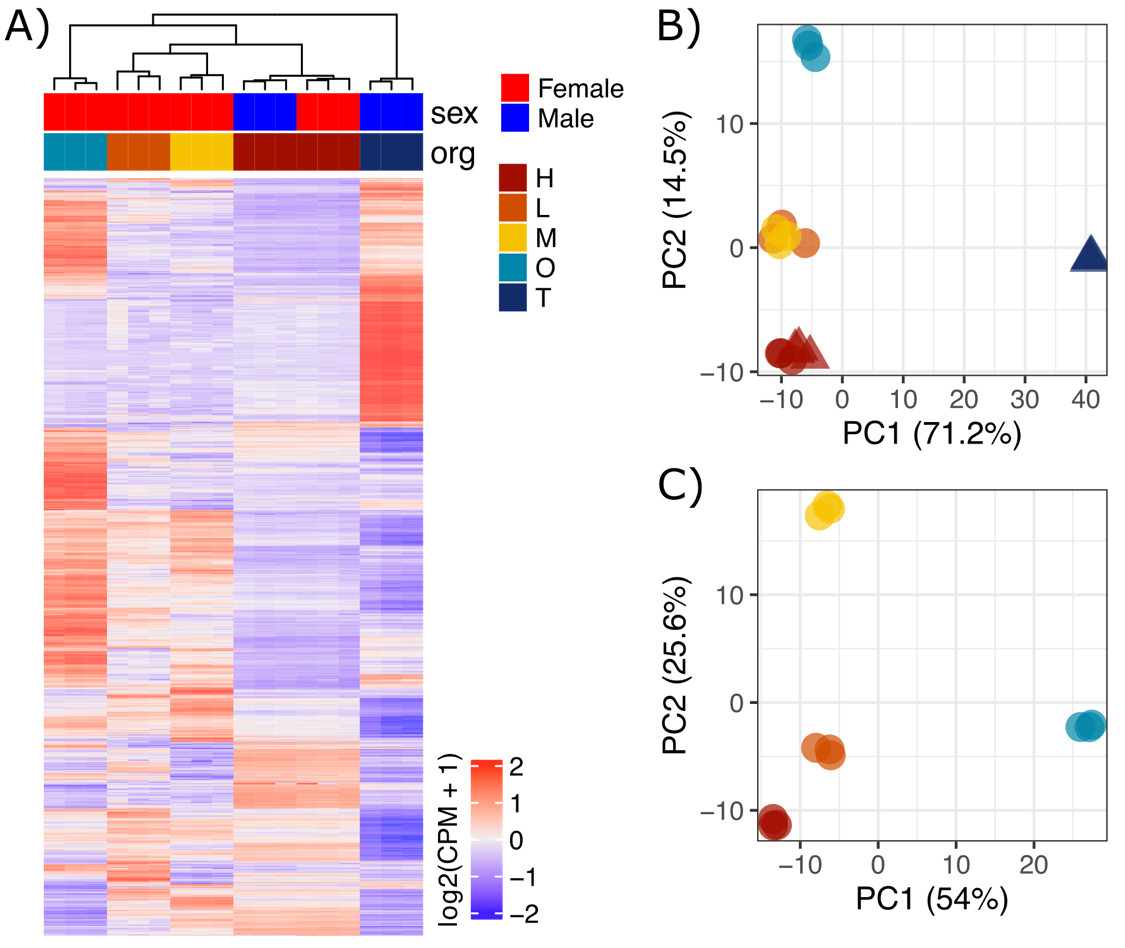

***Figure S2.*** ***A)*** *Heatmap for all genes for all tissues z-transformed log2(cpm + 1).* ***B)*** *Principal component analysis for 500 most variable genes including male heads and testis.* ***C)*** *Principal component analysis for 500 most variable genes for female tissues only.*

Recalculating tissue-specificity combining mesospermalege and lower reproductive tract showed that an additional 277 genes have specific expression across the female reproductive system. As expected, the total combined female reproductive system specific genes include the majority of mesospermalege and lower reproductive tract specific genes (Fig. S3).

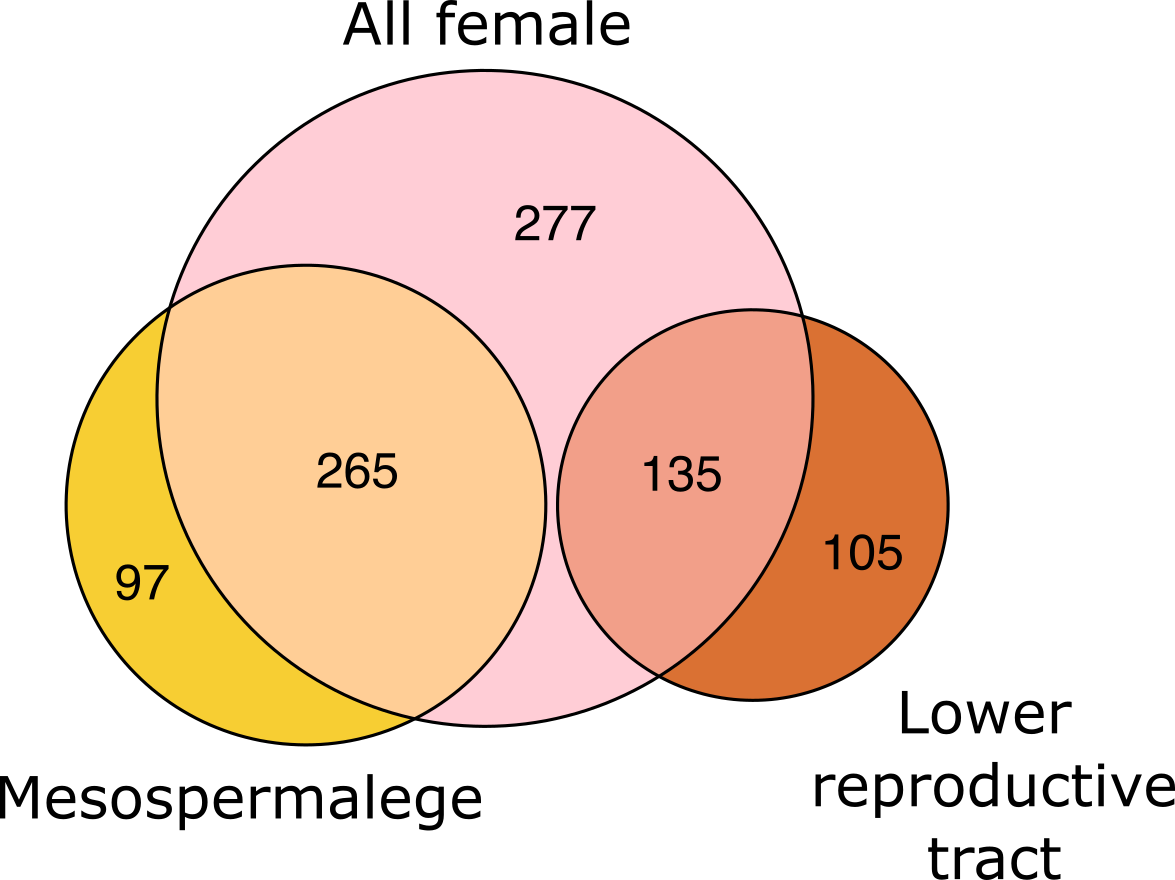

***Figure S3.*** *Overlap of tissue-specific genes in the mesospermalege and lower reproductive tract compared to tissue-specificity calculated for a combined female reproductive tissue class – combining the two female tissues.*

***Chromosomal distribution***

Lower reproductive tract-specific genes were enriched on chromosome 4 ($X^{2}$ = 18.1, df = 1, *p* < 0.001) and ovary-specific genes were enriched on X_1_ ($X^{2}$ = 40.1, df = 1, *p* < 0.001) and X_2_ ($X^{2}$ = 15.4, df = 1, *p* < 0.001) and under-represented on chromosome 10 ($X^{2}$ = 17.3, df = 1, *p* < 0.001; Fig. S4).

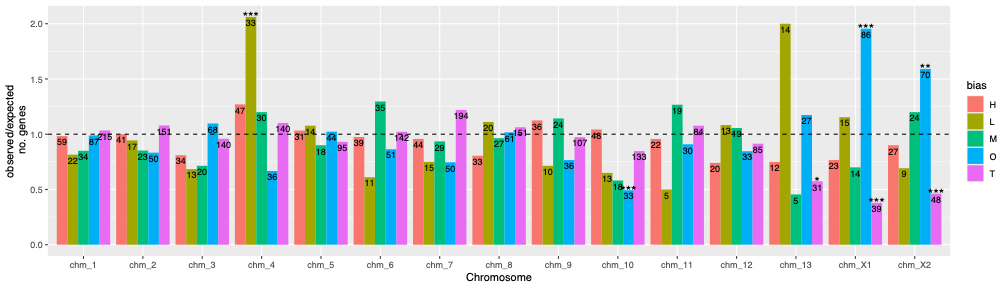

***Figure S4.*** *Chromosomal distribution of female reproductive tract genes, heads, and testes. The dashed line indicates the null expectation and numbers within bars are the numbers of observed genes. Asterisks represent results from comparing the observed to expected number of genes after multiple testing correction using* $X^{2}$ *tests: **: p < 0.01; ***: p < 0.001.*

***Expression of* Drosophila *orthologs***

We identified a number of *D. melanogaster* orthologs in female bedbug tissues. The *sex peptide receptor* (*SPR*) did not pass our filtering step in the main analysis (≥ 5 cpm in ≥ 3 replicates) but showed highest expression in the mesospermalege using the unfiltered TPM data with TPM ≥ 1 (Fig. S5).

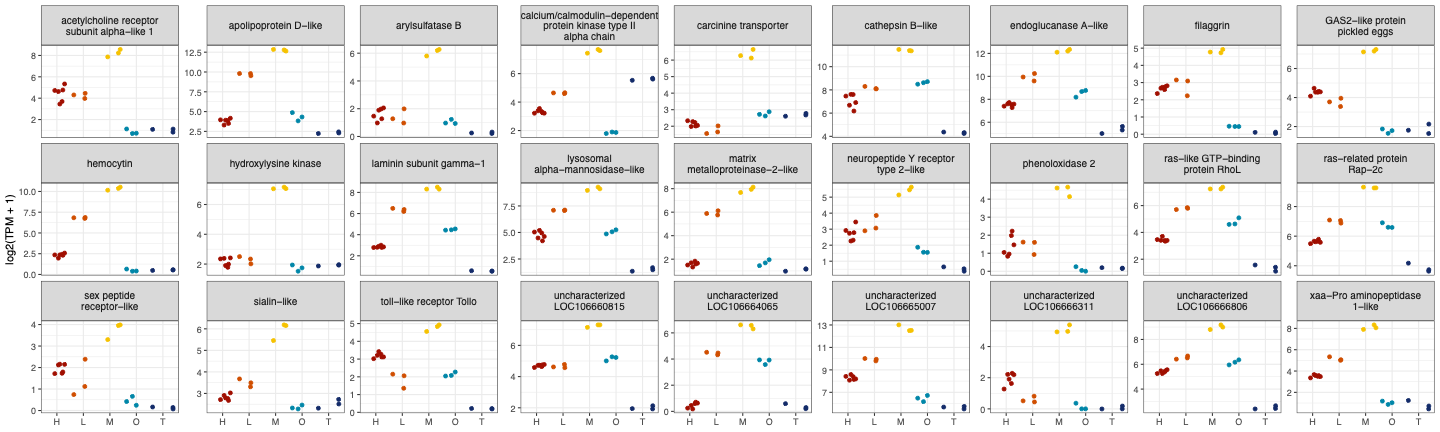

***Figure S5.*** *Expression (transcripts per million reads, TPM) of bedbug orthologs of*

Drosophila melanogaster *genes with highest expression in the mesospermalege.*

In the mesospermalege, the first four PCs explained more than 81% of variance in gene expression across timepoints. PC1 (38.5% variance explained) and PC2 (29.3% variance explained) separated earlier from later timepoints (3h, 6h, and 24h). Samples collected immediately after mating (0h) overlapped with unmated samples (Fig. S6A). In the lower reproductive tract, the first two PCs explain more than 49% of variance in gene expression (Fig. S6B). PC1 (34.9% variance explained) and PC2 (14.5% variance explained) separate the unmated and earlier timepoints (0h and 3h) from the two later time points (6h and 24h).

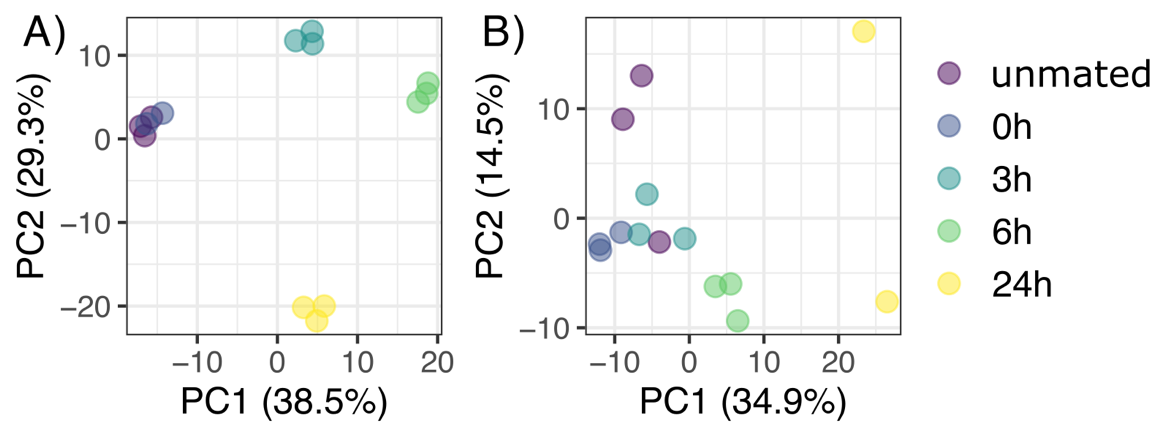

***Figure S6.*** *Principal component analysis for the 500 most variable genes across timepoints in* ***A)*** *the mesospermalege and* ***B)*** *lower reproductive tract.*

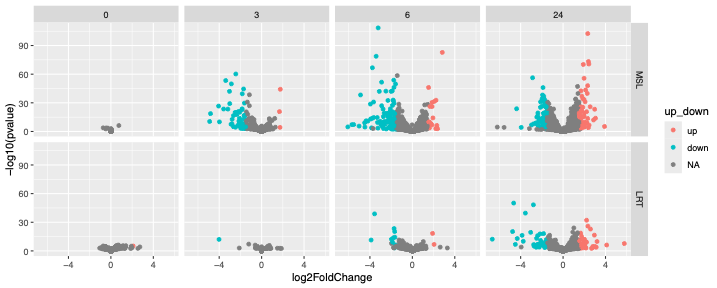

***Figure S7.*** *Volcano plots showing genes differentially expressed compared to unmated samples in the mesospermalege (top) and lower reproductive tract (bottom). Differentially expressed genes (log2FC > |1.5| & adjusted p-value ≤ 0.05) are coloured by higher (red) or lower (blue) expression compared to unmated samples.*

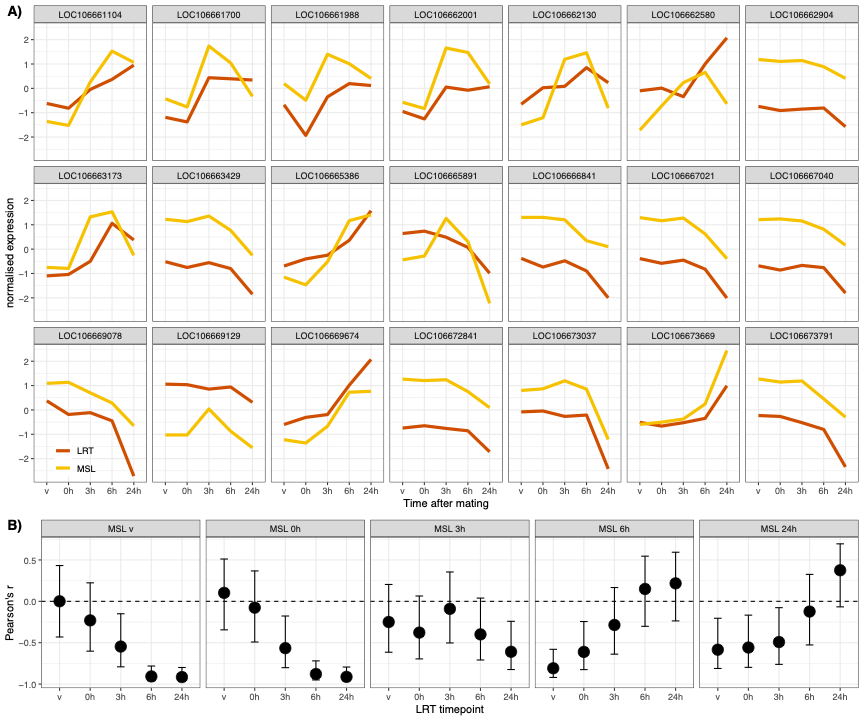

***Figure S8.*** *Co-expression of genes showing postmating transcriptional response in the mesospermalege and lower reproductive tract (n = 21).* ***A)*** *Expression profile for each gene in the mesospermalege (yellow) and lower reproductive tract (orange).* ***B)*** *Pearson’s correlation coefficients (± 95% confidence interval) comparing overall gene expression in the mesospermalege vs. lower reproductive tract at each timepoint.*
